# WikiMedMap: Expanding the Phenotyping Mapping Toolbox Using Wikipedia

**DOI:** 10.1101/727792

**Authors:** Lina Sulieman, Patrick Wu, Joshua C. Denny, Lisa Bastarache

## Abstract

Researchers utilizing phenotypic data from diverse sources require matching of phenotypes to standard clinical vocabularies. Mapping phenotypes to vocabulary can be difficult, as existing tools are often incomplete, can be difficult to access, and can be cumbersome to use, especially for non-experts. We created WikiMedMap as a simple tool that leverages Wikipedia and maps phenotype strings to standard clinical vocabularies. We assessed WikiMedMap by mapping phenotype strings from questionnaires in the UK Biobank and from Mendelian diseases in Online Mendelian Inheritance in Man (OMIM) database to eight vocabularies: International Classification of Diseases, Ninth Revision (ICD-9), ICD-10, ICD-O, Medical Subject Headings (MeSH), OMIM, Disease Database, and MedlinePlus. WikiMedMap outperformed conventional mapping tools in finding potential matches for phenotype strings. We envision WikiMedMap as a technique that complements existing and established tools to map strings to clinical vocabularies that usually do not coexist in one source.

## Introduction

With the completion of the Human Genome Project in 2000, many genome-wide association studies (GWAS) have been published with the goal of understanding how the genetic variation influences phenotypes, including diseases. In the past decade, researchers have used increasingly large datasets and meta analyses to conduct genotype-phenotype association studies at much larger scales. Large cohorts like the UK Biobank (UKBB) (Sudlow et al., 2015), Million Veteran Program (Gaziano et al., 2016), and *All of Us* Research Program (Collins and Varmus, 2015) include dense phenotypic information from a variety of sources, including healthcare data, participant surveys, and objective measurements from in-person visits. Efforts such as the Electronic Medical Records and Genomics (eMERGE) Network (McCarty et al., 2011) and the Million Veteran Program (Gaziano et al., 2016) have demonstrated the power of electronic health records (EHRs) for genomic research. In addition, resources like the genomewide association study (GWAS) catalog (Buniello et al., 2019) have curated data from thousands of studies for a diverse set of diseases and traits. Initiatives like the Undiagnosed Disease Network (Splinter et al., 2018) and MatchMaker (Buske et al., 2015) have developed large datasets of dense phenotypic information for rare diseases. Though “big data” has the potential to improve biomedical research, it can also create unique challenges in the biomedical research space.

Despite the availability of these datasets, it is often challenging to use phenotypic data to answer questions in biomedicine. While there are widely accepted standards for referencing genetic variants, (1000 Genomes Project Consortium et al., 2015; Kent et al., 2002) there are no universally accepted standards or guidelines for describing the human phenome (Banda et al., 2018; Wilcox, 2015). To describe human phenotypes, researchers have developed many ontologies and vocabularies to meet the needs of particular domains and use cases (Petersen et al., 1999; Shivade et al., 2014). For example, the World Health Organization developed International Classifications of Diseases (ICD) codes to track mortality and morbidity, and is commonly used for diagnosis and billing in healthcare (Steindel, 2010). A second example is National Library of Medicine’s (NLM) Medical Subject Headings (MeSH) vocabulary that is used to index scientific publications (Lipscomb, 2000). Third, Human Phenotype Ontology (HPO) was developed to describe the signs and symptoms of Mendelian disease (Robinson et al., 2008). Fourth, Online Mendelian Inheritance in Man (OMIM) and Orphanet were developed to specify monogenic conditions (McKusick, 2007). And fifth, Systematized Nomenclature of Medicine (SNOMED) is a standard terminology commonly used to represent diseases (Cornet and de Keizer, 2008).

Research groups have leveraged these disparate resources to create specific disease cohorts using EHR data by creating phenotype algorithms that map ICD diagnosis codes and other elements to phenotypes of interest (Denny et al., 2013, 2010; Hripcsak et al., 2018; Kirby et al., 2016; Newton et al., 2013; Wu et al., 2019). However, variability in the description of human phenotypes can impede researchers from efficiently conducting genotype-phenotype association studies (Sabb et al., 2009). For example, a researcher studying ovarian cancer in the UKBB must first identify the relevant data elements, which might be ICD10-CM codes or text in the survey questionnaires. At this time, this process cannot be fully automated, as there are many synonyms for “ovarian cancer”, including “ovarian carcinoma” and “Malignant neoplasm of ovary” (ICD-10 C56). To find an equivalent code for “ovarian cancer”, a researcher who is inexperienced phenotyping can review the literature or collaborate with clinical colleagues. While a purely manual effort is manageable for a small number of phenotypes, it is hard to scale this process for hundreds to thousands of phenotypes. Further, these linking efforts are continuous, as new phenotype maps are required when clinical vocabularies are updated and new sources of phenotype information are made available. Hence, tools that partially automate the linking of phenotypes from disparate data sources to standard vocabularies are essential.

Some current tools and databases that facilitate translation between different medical and scientific vocabularies have allowed genotype-phenotype association studies to be conducted using EHR data. However, researchers may not have access to all the vocabularies or have the training necessary to use them appropriately. For instance, researchers commonly use the Unified Medical Language System (UMLS) (Humphreys et al., 1998), a medical terminology thesaurus that binds terms from >100 medical vocabularies to unique concept identifiers and thus can serve as an interlingua between many of these vocabularies. However, the UMLS is not a comprehensive solution to translate across vocabularies. It can be difficult to use, may not include some clinical vocabularies (e.g., consumer health vocabularies or cohort questionnaires), and may lack the necessary synonymy and some relationships (Geller et al., 2009; Zeng and Tse, 2006). The complex data structures and concept relationships in these resources creates an obstacle for scientists interested in mapping their phenotypes to a standard vocabulary, which can impede their research. Even for researchers who have extensive experience working with UMLS, automated mapping often fails to identify equivalent standard medical concepts and require input from domain experts familiar with both the phenotype in question and the standard clinical terminology (Hripcsak et al., 2018).

In this paper, we present a Wikipedia-based tool called WikiMedMap, an open-source tool designed to facilitate the translation of phenotype strings to standard vocabularies. WikiMedMap takes raw strings as inputs and outputs standard codes from common medical vocabularies including ICD-9, ICD-10, MeSH, DiseaseDB, OMIM, and Orphanet. Wikipedia has three features that make it a useful resource for linking medical terms. First, it has a large database of alternative names, common misspellings, and abbreviations to redirect search strings to the correct page (e.g. “regional enteritis” is redirected to the page “Crohn’s disease”), which is a powerful resource for string normalization. Second, pages for medical concepts in Wikipedia are structured using a specially-designed template that includes an “infobox” with classifications using a set of standard terminologies. Third, Wikipedia has an API to execute searches and access results. While it is not comprehensive, WikiMedMap is easy to use for identifying relationships between terms that may not be present in conventional resources like the UMLS.

## Results

### Comparison of WikiMedMap to UMLS

To assess the potential of WikiMedMap in the phenotyping pipeline, we used 691 UKBB phenotypes strings that UKBB participants were asked in questionnaires, to compare WikiMedMap with UMLS. We defined a match as a phenotype with at least one link between the phenotype and the standard medical vocabulary in question. Overall, we found that WikiMedMap finds more standard concept links than UMLS for UKBB terms for all tested vocabularies (Tables 1 and 2). For example, WikiMedMap found ICD-9 codes for 33 UKBB disease terms compared to 57 found with the UMLS. We also compared the performance of WikiMedMap to UMLS in mapping 347 HPO terms (Table 2). WikiMedMap found more links between the HPO terms for 0.75 (6/8) of the standard clinical vocabularies. UMLS found more concept links in MeSH (WikiMedMap 153, UMLS 170) and OMIM (WikiMedMap 71, UMLS 253).

**Table 1.**
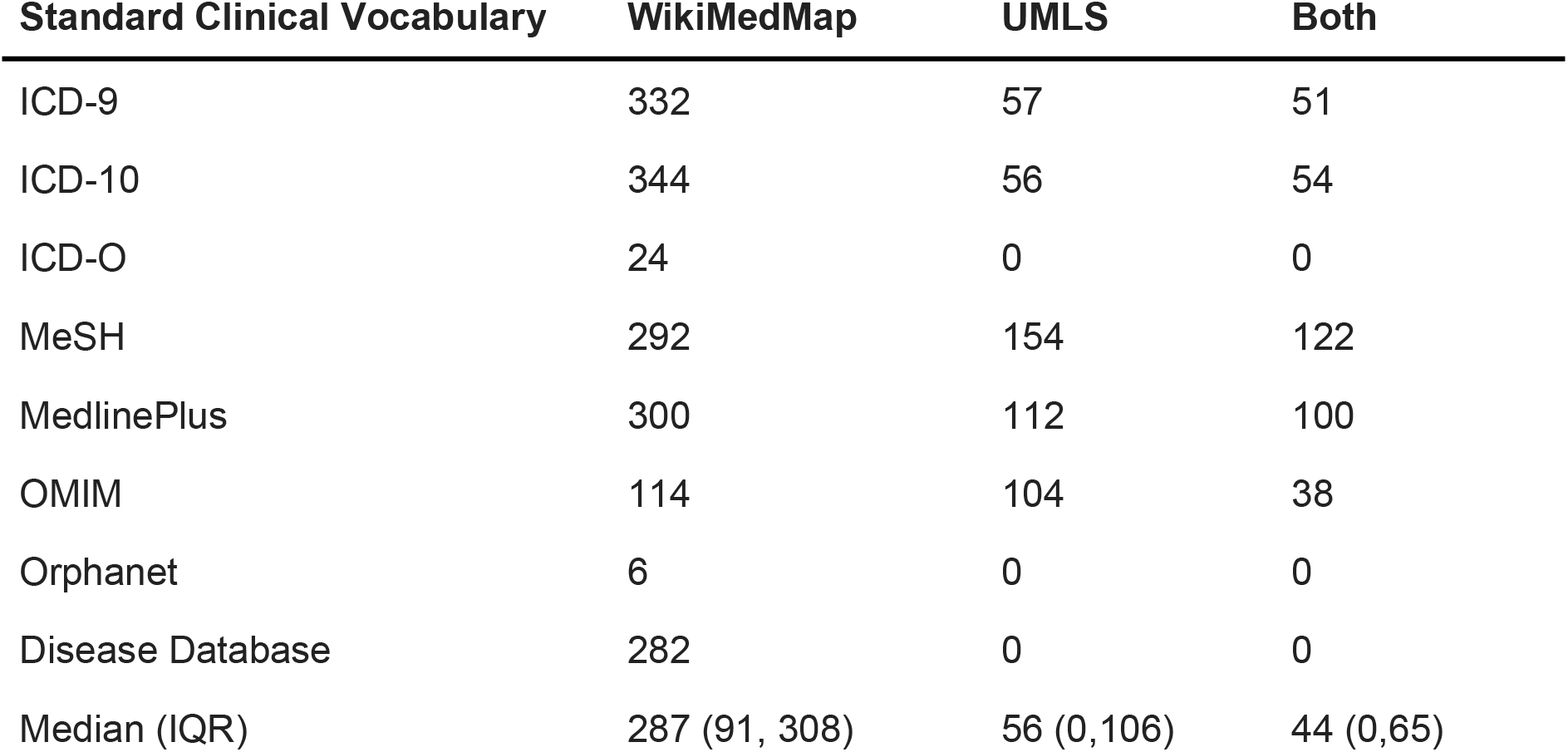
Number of UKBB phenotypes with ≥1 mappings for each standard clinical vocabulary for the 691 UKBB phenotypes.

**Table 2.**
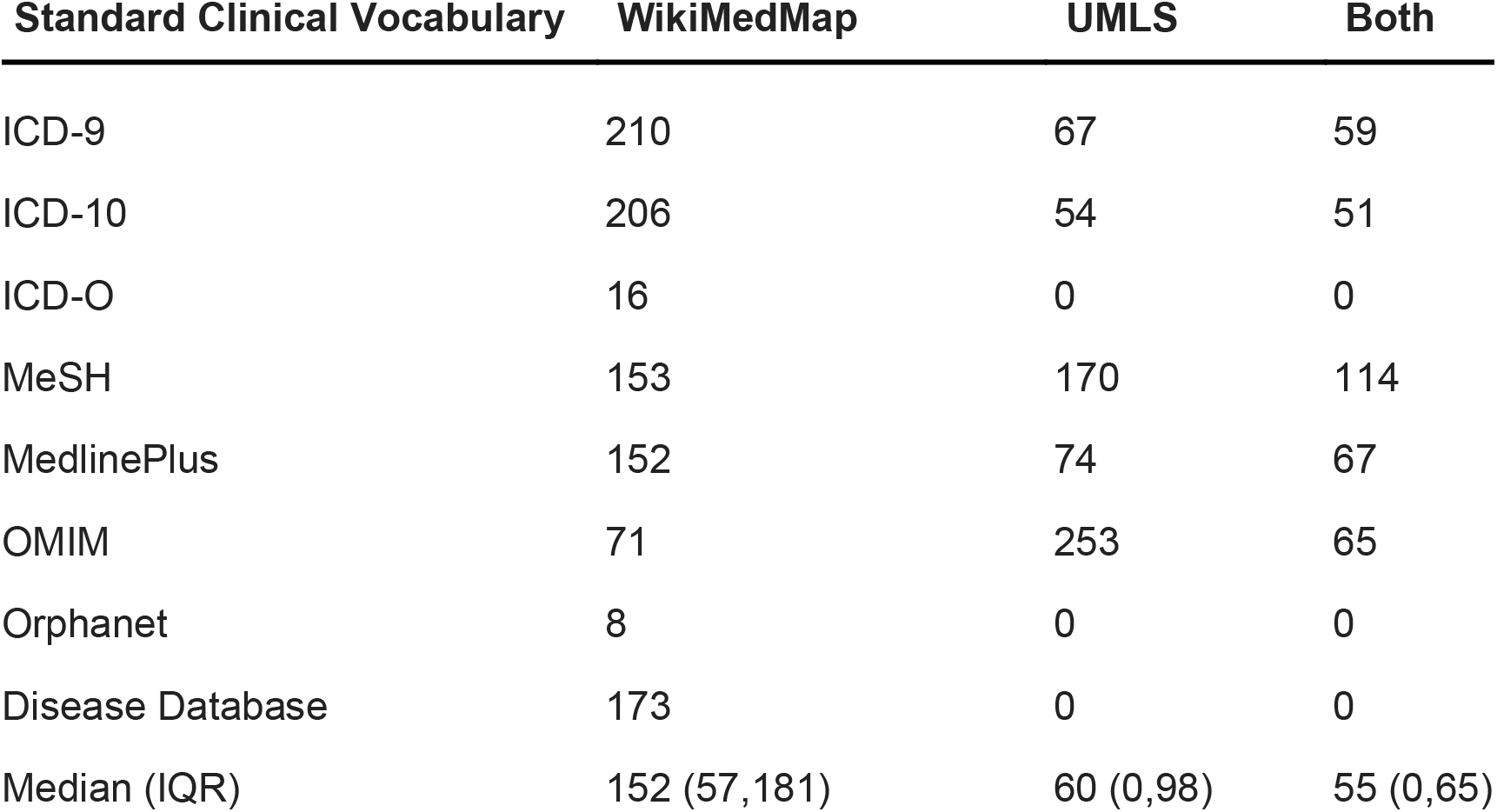
Number of HPO phenotypes with ≥1 mappings for each standard clinical vocabulary for 347 HPO terms

| Standard Clinical Vocabulary | WikiMedMap | UMLS | Both |
| --- | --- | --- | --- |
| ICD-9 | 210 | 67 | 59 |
| ICD-10 | 206 | 54 | 51 |
| ICD-O | 16 | 0 | 0 |
| MeSH | 153 | 170 | 114 |
| MedlinePlus | 152 | 74 | 67 |
| OMIM | 71 | 253 | 65 |
| Orphanet | 8 | 0 | 0 |
| Disease Database | 173 | 0 | 0 |
| Median (IQR) | 152 (57,181) | 60 (0,98) | 55 (0,65) |

To evaluate the performance of WikiMedMap in mapping phenotypes to standard clinical vocabulary, we compared the ICD-9 mappings from UMLS to those found using WikiMedMap. Compared to UMLS mapping of UKBB terms, WikiMedMap successfully mapped the same 0.89 (51/57) of terms to ICD-9, 0.96 (54/56) of terms to ICD-10, 0.79 (122/154) of terms to MeSH terms, 0.89 (100/112) of terms to MedlinePlus, and 0.37 (38/104) of terms to OMIM. For HPO terms, WikiMedMap and UMLS mapped 0.88 (59/67), 0.94 (51/54), 0.67 (114/170), 0.91 (67/74), and 0.26 (65/253) of the same terms to ICD-9, ICD-10, MeSH, MedlinePlus, and OMIM respectively.

## Discussion

In this study, we developed and applied WikiMedMap, a tool that mines Wikipedia to match phenotypes strings to standard clinical vocabularies. Compared to UMLS, WikiMedMap found matching clinical vocabulary codes for more UKBB phenotypes and HPO terms. We envision WikiMedMap as a complementary tool to existing resources by matching concepts codes for phenotype strings that may not be found with automated mapping strategies using conventional databases. By facilitating the mapping of phenotypes to standard clinical vocabularies, WikiMedMap can resolve a key bottleneck in biomedical research and has the potential to accelerate the translation of biomedical data to improve human health. We do not envision researchers with extensive training in biomedical data science and/or are familiar with UMLS and its derivatives as the primary users of WikiMedMap. On the other hand, biomedical researchers who are unfamiliar with biomedical data science tools, but are interested in conducting genotype-phenotype association studies can benefit from this tool. For example, WikiMedMap may be useful for researchers interested in extracting medical concepts from patient survey questionnaires, consumer health data, and disease community forums.

Since WikiMedMap leverages Wikipedia’s web search tool, it has the built-in ability to handle disease acronyms, misspellings, and synonyms. Healthcare providers commonly use acronyms in clinical notes to save time in documenting patient interactions. In some cases, disease acronyms in clinical notes that are unfamiliar to researchers may impede phenotype mapping. For example, “Diastolic heart failure”, “diastolic dysfunction”, and “Heart Failure with preserved Ejection Fraction” refers to the evolving nomenclature of the same clinical entity and is commonly abbreviated “HFpEF”. For this phenotype, WikiMedMap redirects “HFpEF” to the correct Wikipedia article (“Heart failure with preserved ejection fraction,” n.d.) and extracts the appropriate codes. Narrative text on the internet, such as those found in disease forums or on Twitter, are other unconventional sources for health data. In those sources, users often describe their symptoms and diseases using lay medical terms (Doing-Harris and Zeng-Treitler, 2011; Zeng and Tse, 2006). WikiMedMap can help researchers map disease names and symptoms in social media posts to standardized medical concept codes. For example, using WikiMedMap to search for “stomach flu” directs to the Wikipedia page for “Gastroenteritis” and extracts the ICD-9 codes 008.8 “Viral enteritis NOS”, 009.0 “Infectious colitis, enteritis, and gastroenteritis”, 009.1 “Colitis, enteritis, and gastroenteritis of presumed infectious origin”, and 558 “Other and unspecified noninfectious gastroenteritis and colitis”.

WikiMedMap can be used to extract codes for phenotypes strings in studies that link genotypes to phenotypes, especially for researchers who are unfamiliar with conventional phenotyping approaches. Researchers may use WikiMedMap to map phenotypes to standard medical codes that traditional mapping methods may miss (e.g., linking “Self-harm” to ICD-10 codes X60-X84), and to map HPO and Mendelian diseases terms to codes in disparate terminologies. Moreover, this tool can empower researchers in their first-pass of hypothesis generation step by providing the codes that help them in identifying the cohort and extracting their EHR data.

We envision that WikiMedMap will have other potential valuable uses for biomedical researchers across numerous domains. Researchers can automatically extract the concepts codes for survey questions or answers rather than following a manual approach. In addition, WikiMedMap may be helpful for creating maps for disease or phenotype terms in languages other than English. We present four potential use cases for WikiMedMap (Figure 1): synonym, medical acronyms, phenotypes with difficulty in identifying equivalents in UMLS, and translation of medical concepts in English to other languages.

**Figure 1.**
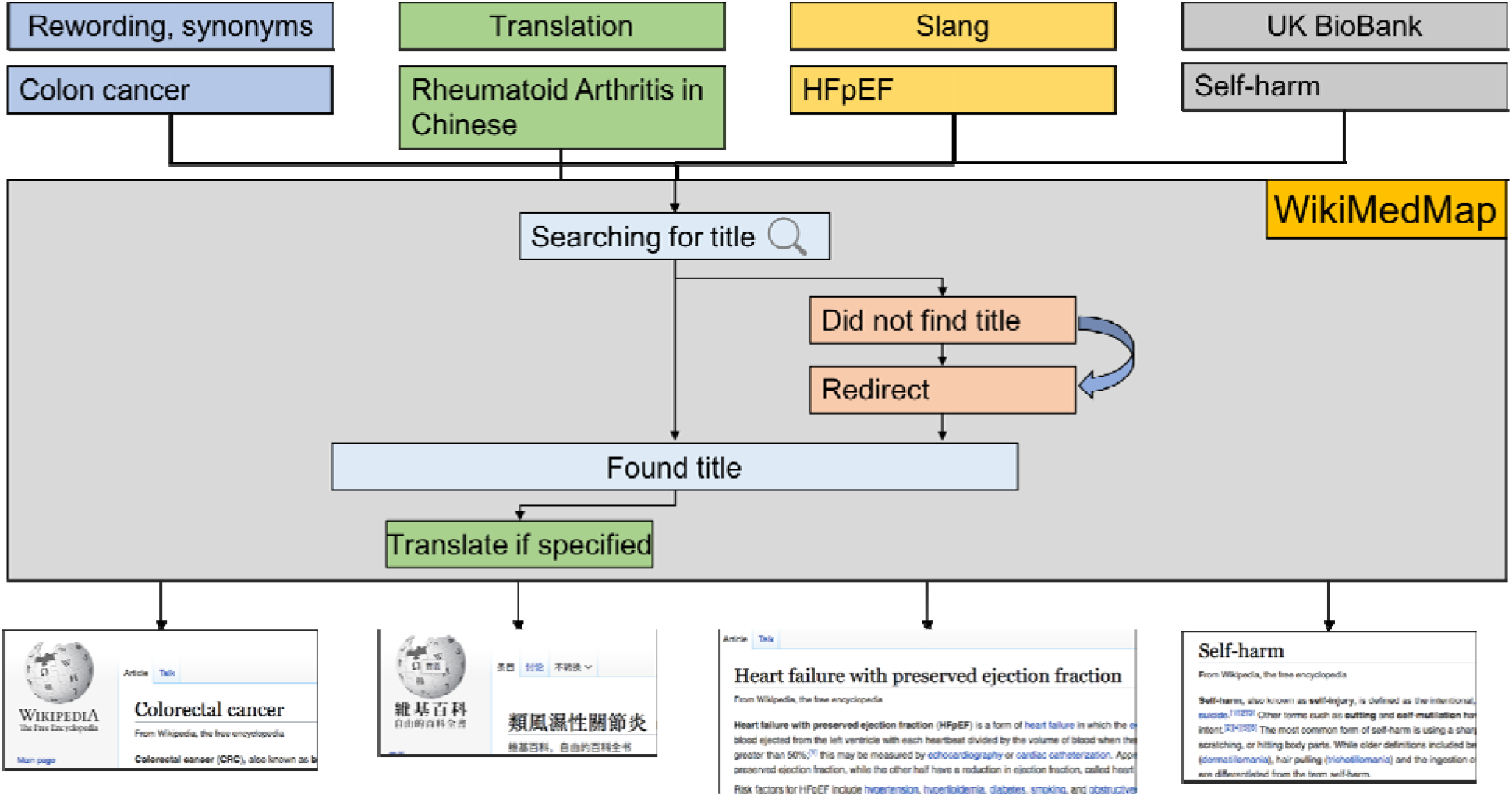
Use cases and extensions of WikiMedMap. For the string ‘colon/cancer’, WikiMedMap identifies the Wikipedia article for ‘Colorectal cancer’ (Wikipedia contributors, 2019) and extracts the relevant ICD codes: ICD-9 153.0-154.1, ICD-10 C18, C19, C20. With WikiMedMap, “Rheumatoid Arthritis” finds the relevant article in the Chinese version of Wikipedia and displays summary from the article to the user. WikiMedMap can understand medical acronyms, such as “Stomach flu”, which redirects to “Gastroenteritis” and extracts ICD-9. WikiMedMap identifies relevant ICD-10 X60-X84 codes that are relevant for the string “self-harm”.

## Limitations

Our tool has some limitations. First, we did not perform a comprehensive analysis to assess the accuracy of WikiMedMap mappings to UMLS. We made this decision, as WikiMedMap is designed to help researchers generate hypotheses, and does not replace standardized vocabularies. We recommend that researchers use more conventional mapping tools or collaborate with individuals with expertise with phenotype mapping for their final studies. Second, though WikiMedMap can parse and analyze disease names from short strings, this tool is not designed to tokenize and analyze clinical notes. For researchers interested in using WikiMedMap to analyze clinical notes, we recommend that researchers pre-process and tokenize clinical notes before feeding the smaller substrings into WikiMedMap. Other more sophisticated natural language processing packages, such as KnowledgeMap (Denny et al., 2003), cTAKES (Savova et al., 2010), and CLAMP (Soysal et al., 2017) can process longer strings and notes. However, they rely on UMLS as the main source for defining the clinical terms which subject them to some of the UMLS limitations discussed earlier. Third, we aimed to simulate real-world scenarios when we compared the performance of WikiMedMap to UMLS. Thus, we used simple string matching instead of concept relationships in the UMLS. We made this decision because we assumed that most of WikiMedMap’s intended users would lack the technical expertise to use this tool in UMLS.

## Future directions

In the future, we plan on expanding this package that has the ability to capture ICD codes ranges. For example, for the string “colon/cancer”, the package currently captures the beginning and end ICD-9 codes, 153.0-154.1. An ideal output for “colon/cancer” would be all the ICD-9 codes between 153.0-154.1. Another potential direction would be to extract more detailed information about phenotypes such as symptoms, complications, and differential diagnosis associated with diseases.

## Conclusions

Biomedical data science tools have allowed researchers to conduct studies that would have been impossible just ten years ago (Ezer and Whitaker, 2019). Phenotype mapping is one bottleneck that impedes translation of biomedical knowledge to improve human health. WikiMedMap facilitates the translation of phenotypes to several commonly used vocabularies and may aid in exploring datasets and generating EHR phenotype algorithms.

## Materials and methods

### WikiMedMap algorithm

WikiMedMap is a standalone package that leverages the Wikipedia API to extract concepts codes from different terminologies documented in Wikipedia. Figure 3 depicts a screenshot of our notebook for the tool. Our tool uses three main components that are easy to install on any computer or cloud service that supports Python. It relies on two python libraries: MediaWiki (“MediaWiki,” n.d.), and Pandas (“Python Data Analysis Library — pandas: Python Data Analysis Library,” n.d.). Pandas is an open source library that supports easy-to-use data structures and high performance computational analysis in Python. MediaWiki is a Python wrapper and parser for the original MediaWiki API, a web service that combines HTML, JSON, and XML to crawl and retrieve Wikipedia articles (“MediaWiki,” n.d.).

**Figure 2.**
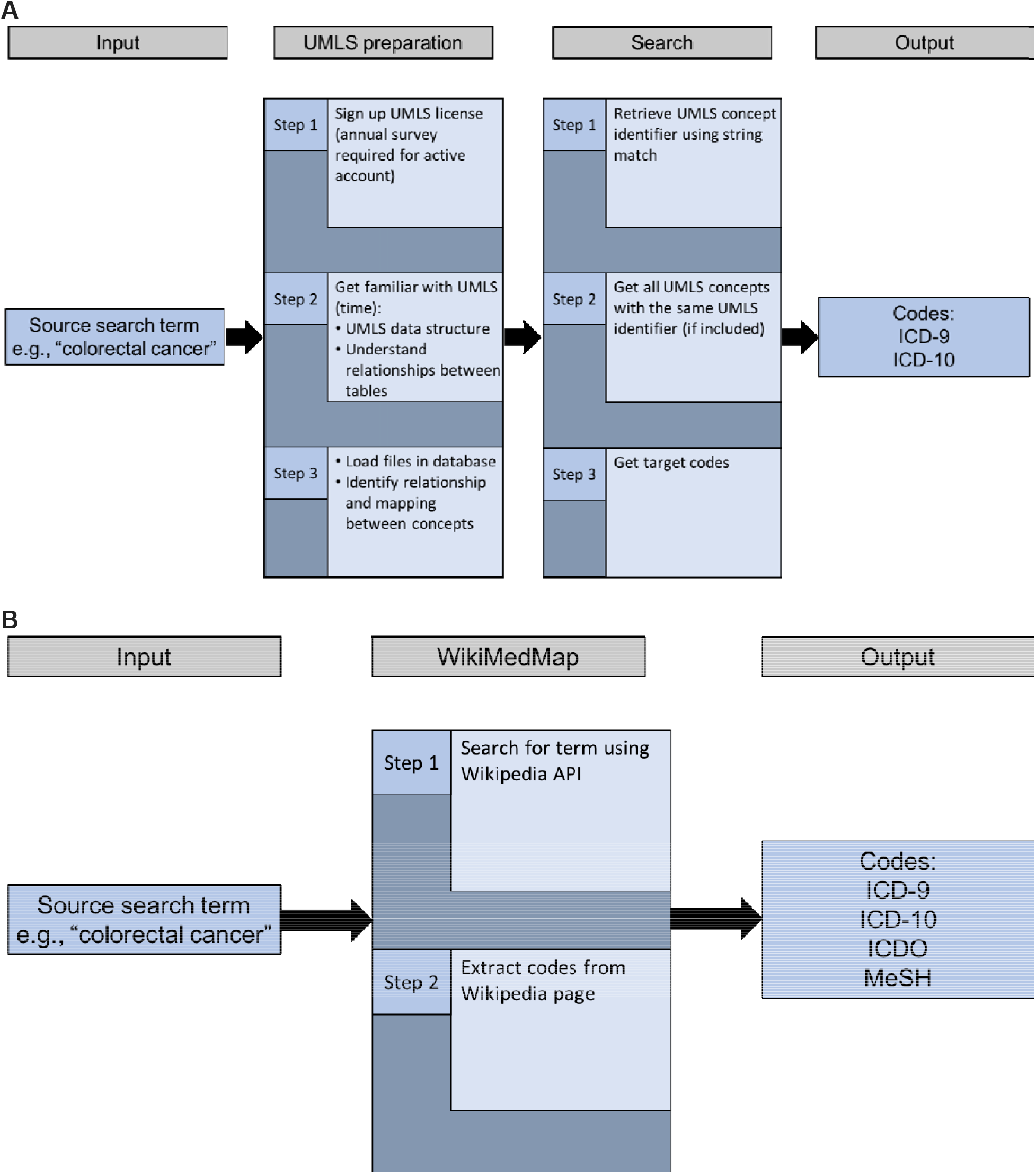
UMLS and WikiMedMap pipeline to map phenotypes. (A) To use UMLS to automate high-throughput mapping of human phenotypes requires many steps. (B) WikiMedMap is simple and robust for automated high-throughput mapping of human phenotypes. UMLS: Unified Medical Language System. ICD: International Classification of Diseases.

**Figure 3.**
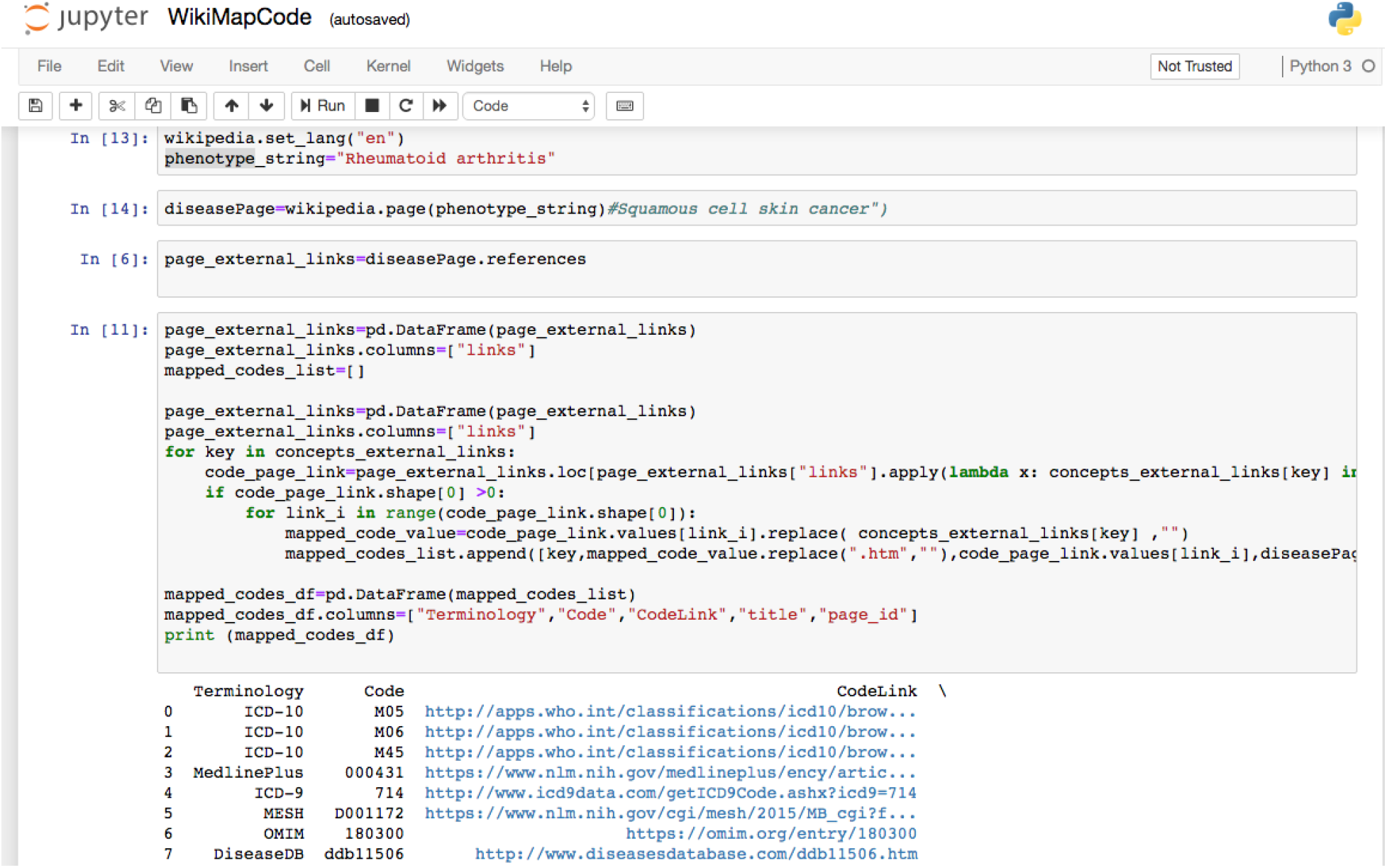
A screenshot of WikiMedMap notebook page

We retrieved the Wikipedia page of the source or the search term using MediaWiki. Then, we parsed the page returned by the API to locate the presence of eight standardized vocabularies: ICD-9, ICD-10, MeSH, MedlinePlus, OMIM, Orphanet, DiseaseDB, and ICD-O. For each terminology, we extracted the codes and the links to the code’s page on the terminology’s website. All analyses were conducted using Python (version 3.6.7), Pandas (version 0.23.4), and Wikipedia API (version 1.4.0).

The Wikipedia API retrieves the most probable page based on the passed source or search string. A user can provide the source string and WikiMedMap returns a list of concept codes from defined vocabularies in the Wikipedia infobox. We used the MediaWiki Python package and located the Wikipedia page of the given term and extracted the page content. MediaWiki returned the most relevant page that represented term whether by exact matching or submatching (Figure 2). Thus, the API often can return the correct Wikipedia entry even if the input string was different from the title of the Wikipedia page. For instance, for “stomach flu”, the API returned the Wikipedia page for “Gastroenteritis”.

### Comparison of WikiMedMap to UMLS

We compared the effectiveness of the WikiMedMap to UMLS in extracting medical vocabularies for strings from two well-known biomedical research repositories: disease substrings from UKBB questionnaires and Mendelian disease terms from OMIM. We extracted codes for each string using UMLS and using WikiMedMap. In UMLS, we used string matching to retrieve vocabulary codes that matched the input string. We avoided using the substring match since doing so may increase the number of false-positive results. We then used the same strings in WikiMedMap, which returned all the matching codes. For UKBB and OMIM strings, we counted the number of unique strings that we were able to successfully map to at least one code for each of the eight standard vocabularies.

## Software and data availability

The WikiMedMap code is available on three platforms: GitHub (https://github.com/Linasulieman/WikiMedMap/blob/master/WikiMedMapCode.ipynb), binder (https://mybinder.org/v2/gh/Linasulieman/WikiMedMap/master), and Google Colab (https://colab.research.google.com/drive/1ZbIPYG6yvM6qRb5Yv8uOUYP38PuIhL2u). The files that include the UKBB surveys and OMIM Mendelian diseases are available as well.

## Acknowledgements

This project was supported by the National Library of Medicine, of the National Institutes of Health, grant R01 LM010685.

